# The projections of ipRGCs and conventional RGCs to retinorecipient brain nuclei

**DOI:** 10.1101/2020.06.17.158022

**Authors:** Corinne Beier, Ze Zhang, Maria Yurgel, Samer Hattar

## Abstract

Retinal ganglion cells (RGCs), the output neurons of the retina, allow us to perceive our visual environment. RGCs respond to rod/cone input through the retinal circuitry, however, a small population of RGCs are in addition intrinsically photosensitive (ipRGCs) and project to unique targets in the brain to modulate a broad range of subconscious visual behaviors such as pupil constriction and circadian photoentrainment. Despite the discovery of ipRGCs nearly two decades ago, there is still little information about how or if conventional RGCs (non-ipRGCs) target ipRGC-recipient nuclei to influence subconscious visual behavior. Using a dual recombinase color strategy, we showed that conventional RGCs innervate many subconscious ipRGC-recipient nuclei, apart from the suprachiasmatic nucleus. We revealed previously unrecognized stratification patterns of retinal innervation from ipRGCs and conventional RGCs in the ventral portion of the lateral geniculate nucleus. Further, we found that the percent innervation of ipRGCs and conventional RGCs across ipsi- and contralateral nuclei differ. Our data provide a blueprint to understand how conventional RGCs and ipRGCs innervate different brain regions to influence subconscious visual behaviors.

## INTRODUCTION

Retinal ganglion cells (RGCs) are the output neurons that connect the eye to the brain and are responsible for all visual perception in mammals. Light information conveyed by RGCs contributes to pattern vision (image forming), as well as subconscious visual behaviors (non-image forming) such as pupil constriction, circadian photoentrainment, sleep and mood. Intrinsically photosensitive RGCs (ipRGCs) are a small population of RGCs that express melanopsin and project to unique targets in the brain that modulate a broad range of non-image forming visual behaviors (Altimus et al., 2008; Chen et al., 2011; Ecker et al., 2010; Hattar et al., 2006; LeGates et al., 2013; Lupi et al., 2008; Rupp et al., 2019). There is still, however, little information about how or if conventional RGCs (non-ipRGCs) can influence subconscious visual behaviors. To address this knowledge gap, we must first delineate whether conventional RGCs innervate non-image forming nuclei.

To determine if conventional RGCs innervate non-image forming nuclei (Ecker et al., 2010; Hattar et al., 2002, 2006; Li and Schmidt, 2018; Morin and Studholme, 2014), we utilized a dual recombinase two color strategy to delineate, in the same animal, how ipRGCs and conventional RGCs converge and diverge in ipRGC-recipient brain regions. We found that the master clock in the suprachiasmatic nucleus (SCN) is exclusively innervated by ipRGCs, as was previously implicated (Hattar et al., 2006). However, the SCN was the exception to the rule, as most ipRGC-recipient areas received substantial projections from conventional RGCs. The dual color strategy allowed us to differentiate areas where conventional RGCs and ipRGCs converge and intermingle, from brain regions where they form completely distinct subdivisions. Specifically, we found that conventional RGCs and ipRGCs innervate distinct subdivisions of the ventral lateral geniculate nucleus (vLGN). As we found in the SCN, the vLGN core region receives exclusive ipRGC input. Furthermore, we found differences in the percent innervation between the ipsi- and contralateral patterns of conventional RGCs and ipRGCs. Our results provide a basis for understanding how ipRGCs and conventional RGC modulate non-image forming behaviors for normal function.

## RESULTS

### The SCN receives exclusive input from ipRGCs

To investigate the relative innervations by conventional RGCs and ipRGCs to retinorecipient nuclei, we developed a method to selectively label both cell populations in a single mouse (Figure 1A). We used the *Opn4*^*Cre*^ mouse line that expresses Cre in ipRGCs, in combination with a Flp-Cre dual recombinase reporter mouse line, RC::FLTG (Rosa26-CAG-Frt-Stop-Frt-loxP-tdTomato-loxP-eGFP, shorthand *Rosa26*^*CAG-tdTomato-eGFP*^) (Ecker et al., 2010; Plummer et al., 2015). In the RC::FLTG mouse line, Flp recombinase excises the Frt flanked stop signal to label Flp-positive cells with tdTomato. The tdTomato transcription termination signal prevents eGFP expression. Cells that are only Cre-positive excise the loxP sites but stay unlabeled while the Frt flanked stop signal remains (Figure 1B). Only cells that express Cre and Flp recombinases are labeled with eGFP (Figure 1C).

**Figure 1.**
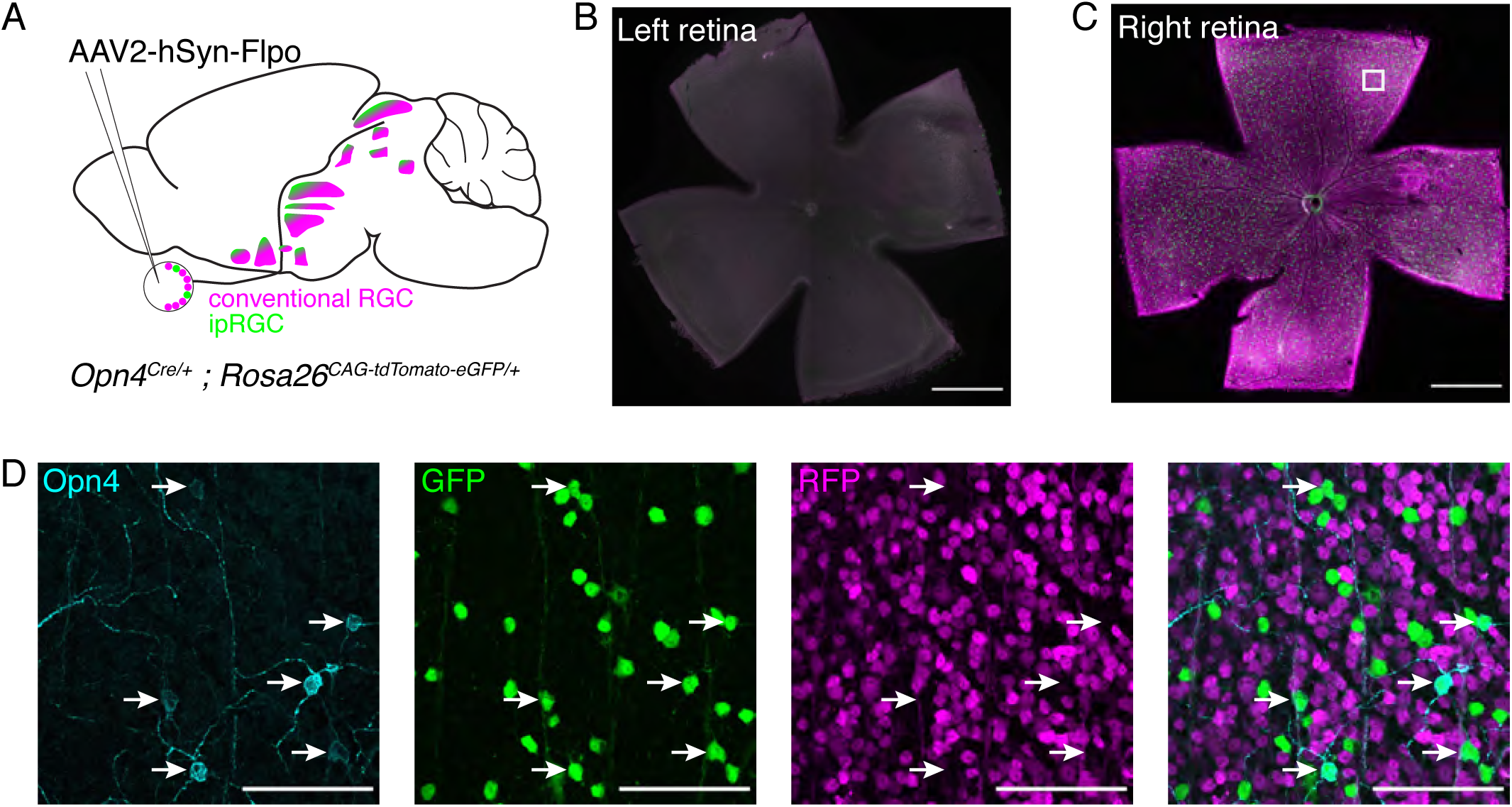
Experiment design and verification. A.Schematic detailing experiment strategy. AAV2-hSyn-Flpo is injected intravitreally into one eye of adult *Opn4*^*Cre/+*^;*Rosa26*^*CAG-tdTomato-eGFP/+*^ mice. Conventional RGC (magenta) and ipRGC (green) projections can be visualized in retinorecipient nuclei. **B**. Immunostaining of GFP and RFP in the retina of the eye without intravitreal injection (left eye) in *Opn4*^*Cre/+*^;*Rosa26*^*CAG-*^ *tdTomato-eGFP/+* mice. No cells are labeled. Scale bar 1000 µm. **C**. Immunostaining of GFP (green) and RFP (magenta) in the retina of the eye that received intravitreal viral injection in *Opn4*^*Cre/+*^;*Rosa26*^*CAG-tdTomato-eGFP/+*^ mice. Scale bar 1000 µm. **D**. Detail view from location indicated in (C). Melanopsin (Opn4) antibody stains a subset of cells (cyan, arrows). All melanopsin-positive cells are GFP-positive (green). RFP-positive cells (magenta) do not overlap with melanopsin- or GFP-positive cells. Scale bar 100 µm.

We generated *Opn4*^*Cre/+*^;*Rosa26*^*CAG-tdTomato-eGFP/+*^ mice and injected AAV2-hSyn-Flpo intravitreally into one eye of adult mice to infect all RGCs with Flp (Figure 1A). AAV2-hSyn-Flpo expresses the Flp recombinase under the neuronal cell specific synapsin promoter. In the retina of *Opn4*^*Cre/+*^;*Rosa26*^*CAG-tdTomato-eGFP/+*^ mice infected with AAV2-hSyn-Flpo, ipRGCs express Flp and Cre, whereas conventional RGCs only express Flp. As expected, this method specifically labels ipRGCs with eGFP and conventional RGCs with tdTomato (Figure 1C). The dual recombinase strategy worked as expected, since no cells expressed both tdTomato and eGFP (Figure 1D). To test whether our strategy is selective to ipRGCs, we stained the retina of *Opn4*^*Cre/+*^;*Rosa26*^*CAG-tdTomato-eGFP/+*^ mice infected with AAV2-hSyn-Flpo with a melanopsin antibody. We found complete overlap between melanopsin expressing cells and eGFP (Figure 1D), confirming that we can selectively label ipRGCs from conventional RGCs.

After confirming our labeling strategy, we traced axonal fibers that project to ipRGC-recipient brain regions to answer the following questions: (1) Do ipRGC-recipient brain regions receive exclusive input from ipRGCs? (2) If not, do the inputs of conventional RGCs and ipRGCs converge or diverge? (3) What are the projection patterns of conventional RGCs and ipRGCs in their ipsilateral and contralateral regions? We focused our analysis on retinorecipient nuclei known to receive robust ipRGC innervation: the suprachiasmatic nucleus (SCN), dorsal lateral geniculate nucleus (dLGN), ventral LGN (vLGN), intergeniculate leaflet (IGL), olivary pretectal nucleus (OPN), and superior colliculus (SC). We also observed modest ipRGC innervation to the zona incerta (ZI), peri habenula (PHb), and supraoptic nucleus (SON). Additional retinorecipient areas that are known to receive minor ipRGC innervation were labeled inconsistently by the dual recombinase mouse and hence were not included in this study (Delwig et al., 2016; Ecker et al., 2010; Hattar et al., 2002, 2006; Li and Schmidt, 2018).

Testifying to the specificity of the dual recombinase strategy, we first examined the SCN, thought to receive exclusive input from ipRGCs (Baver et al., 2008; Chew et al., 2017; Güler et al., 2008; Hattar et al., 2006). We show indeed that the SCN barely receives any input from conventional RGCs (Figure 2, inset in middle panel). From rostral to caudal, the SCN receives only ipRGC innervations that are uniform across the ipsilateral and contralateral regions (Figure 2). The ipsilateral SCN nucleus received 48% of ipRGC innervation and this innervation was equivalent across the ipsi- and contralateral regions (Student’s paired t test, p = 0.31).

**Figure 2.**
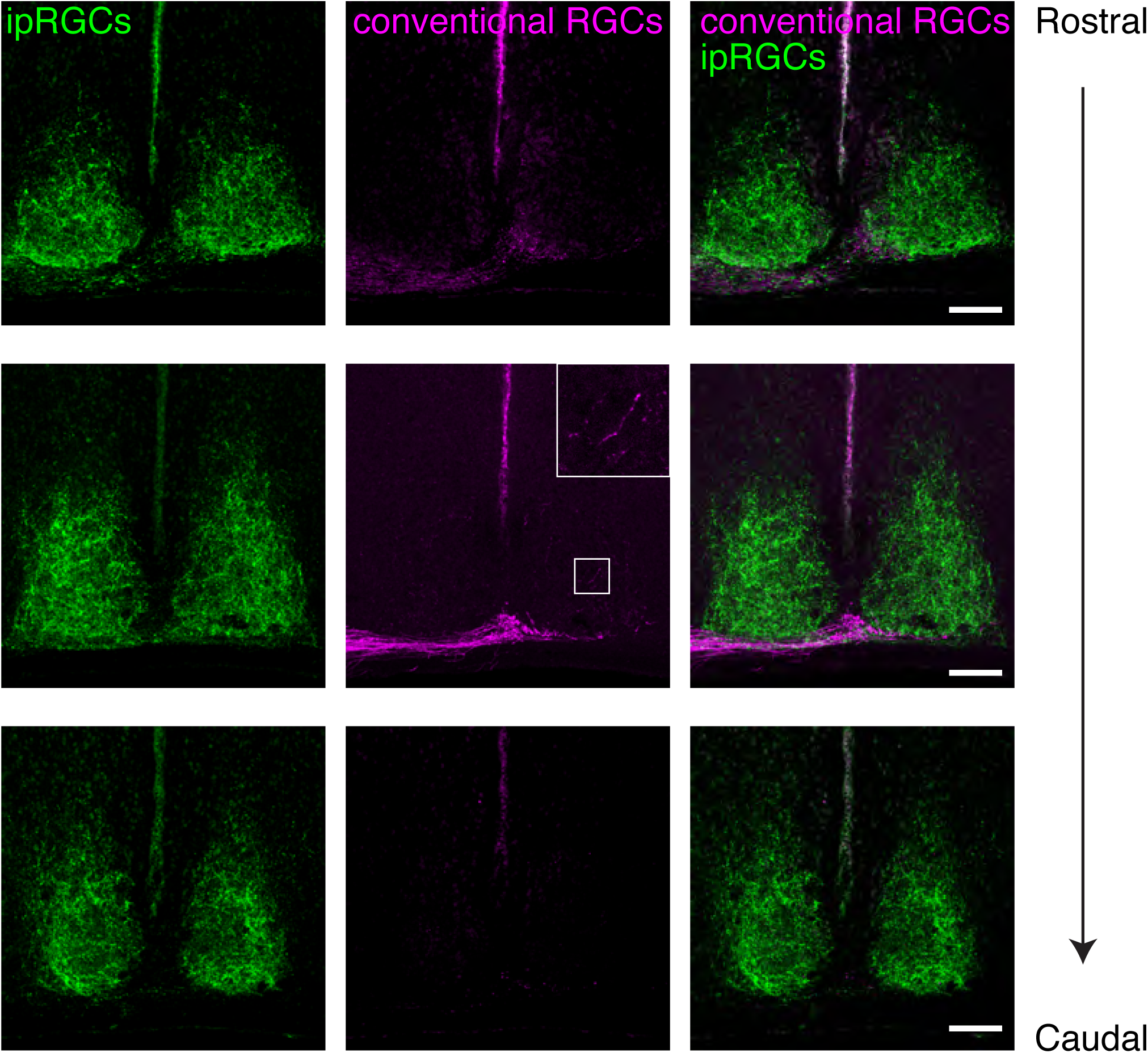
ipRGC and conventional RGC projections in the SCN. ipRGC (green) and conventional RGC (magenta, inset) innervation to the suprachiasmatic nucleus (SCN). ipRGC input is uniform across the contra (left) and ipsilateral (right) lobes. Scale bars 100 µm.

### ipRGC projections overlap with conventional RGCs in the SC and dLGN

A subset of ipRGCs play a role in image-forming vision and project to the image-forming retinorecipient targets, the SC and dLGN (Ecker et al., 2010; Estevez et al., 2012; Quattrochi et al., 2018; Schmidt et al., 2014; Stabio et al., 2017). In the contralateral SC, ipRGC and conventional RGC innervations overlapped in the retinorecipient layer with ipRGC innervation appearing patchy (Figure 3). However, ipRGC innervation was confined to the deeper areas of the SC while conventional RGC innervation covered the entire region (Figure 3). The ipsilateral SC received minimal, but completely overlapping ipRGC and conventional RGC input (Figure 3).

**Figure 3.**
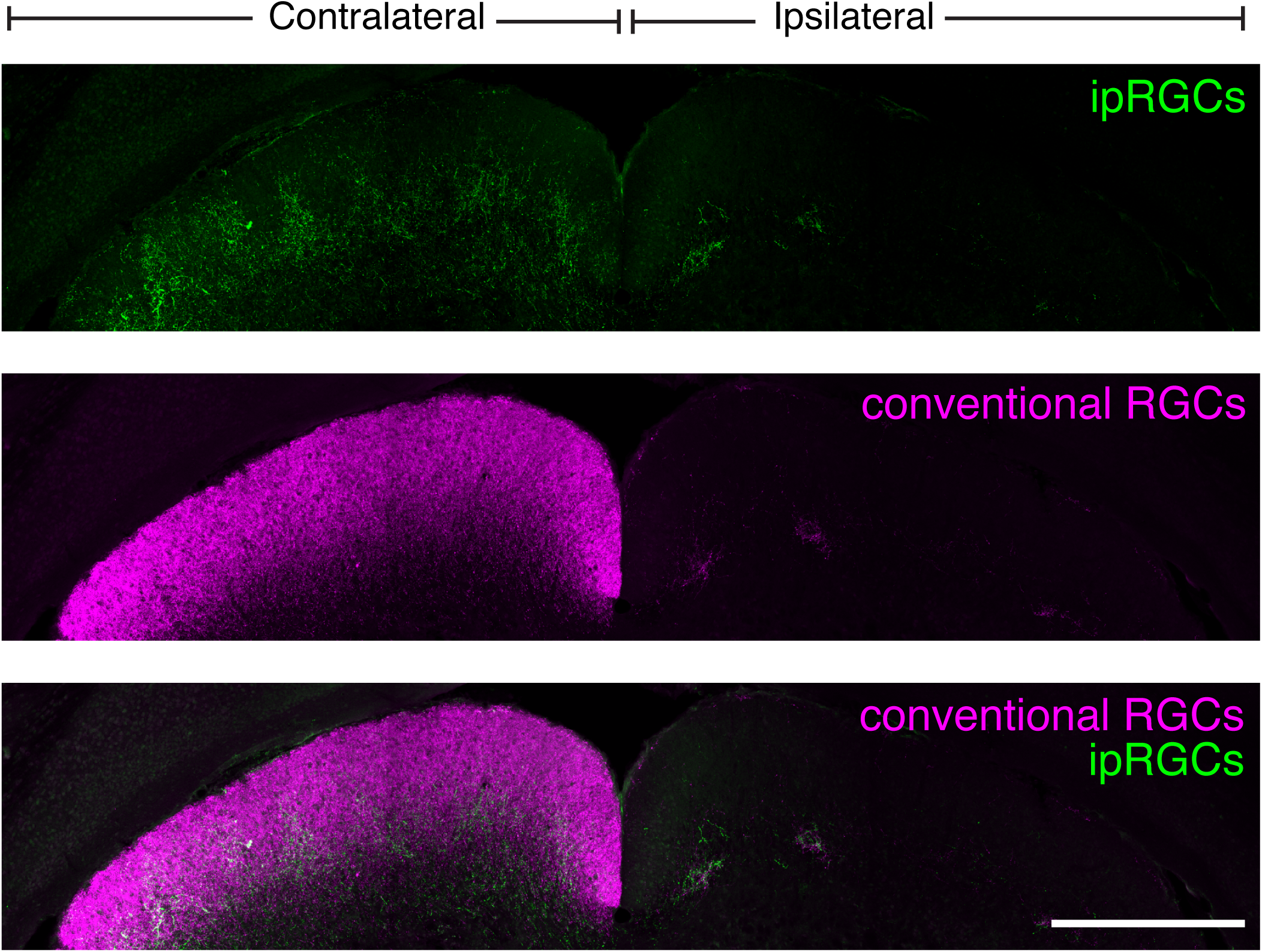
ipRGC and conventional RGC projections in the SC. ipRGC (green) and conventional RGC (magenta) innervation to the contralateral (left) and ipsilateral (right) superior colliculus (SC). Scale bar 500 µm.

In the dLGN, we also observed overlapping ipRGC and conventional RGC innervation (Figure 4). As we have shown previously (Ecker et al., 2010), ipRGC innervation is concentrated towards the medial border of the dLGN nucleus and this pattern becomes more apparent in the caudal dLGN (Figure 4). However, this area was not exclusive to ipRGCs as conventional RGCs innervated the entirety of the dLGN, including the medial border (Figure 4). In the ipsilateral dLGN, there was considerable overlap between the ipRGCs and conventional RGCs (Figure 4), with ipRGCs contributing substantially to the ipsilateral region despite being a minority population in the retina (Figure 5).

**Figure 4.**
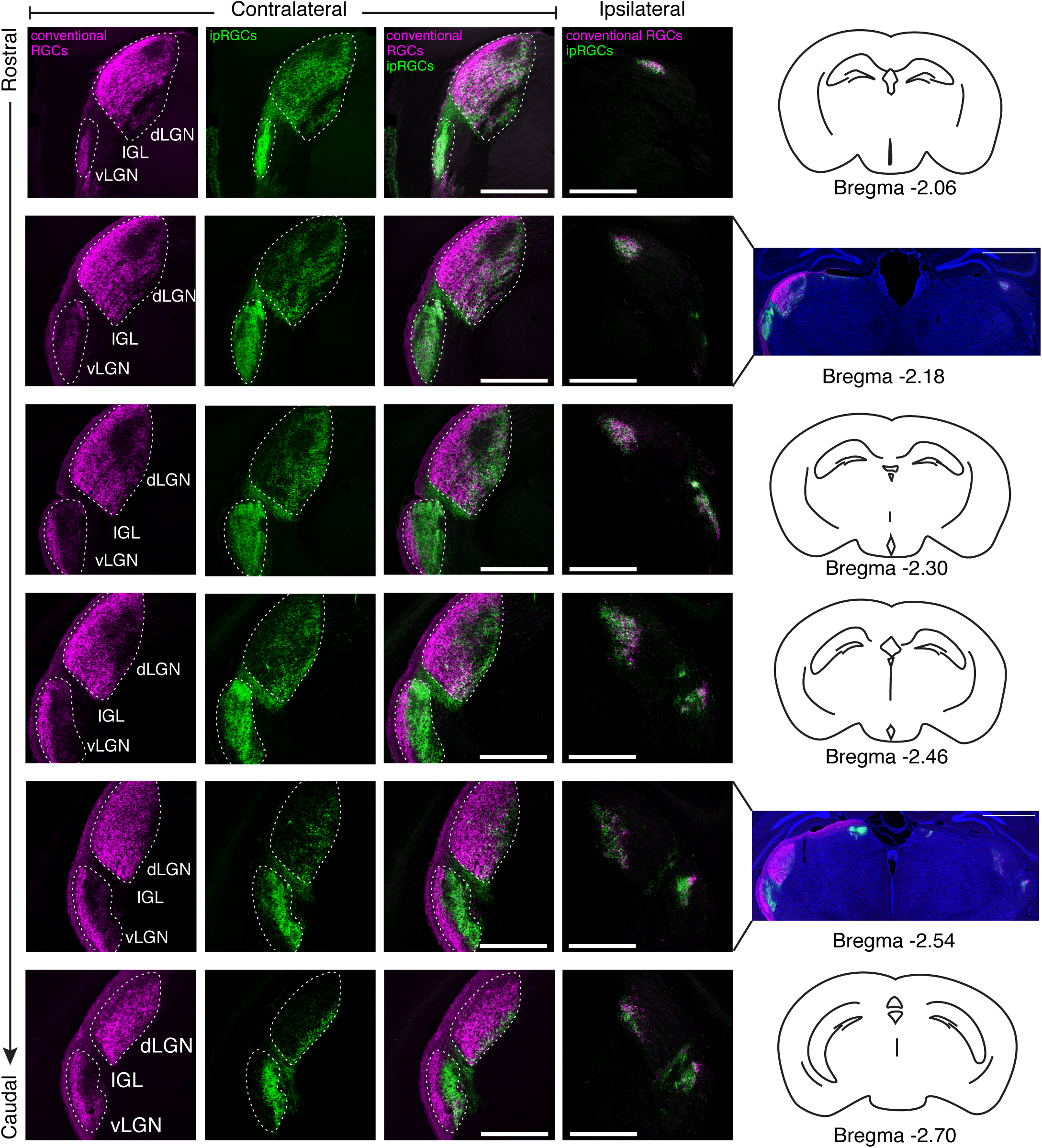
ipRGC and conventional RGC projections to the geniculate complex. Conventional RGC (magenta) and ipRGC (green) innervation to the contralateral and ipsilateral geniculate. The dorsal lateral geniculate nucleus (dLGN) and the ventral LGN (vLGN) are separated by a thin region, the intergeniculate leaflet (IGL). Retinal innervations to the contralateral dLGN and vLGN are outlined (dashed lines). Scale bars 500 µm. Sections are displayed top to bottom, from rostral to caudal, with distances from Bregma listed far right.

**Figure 5.**
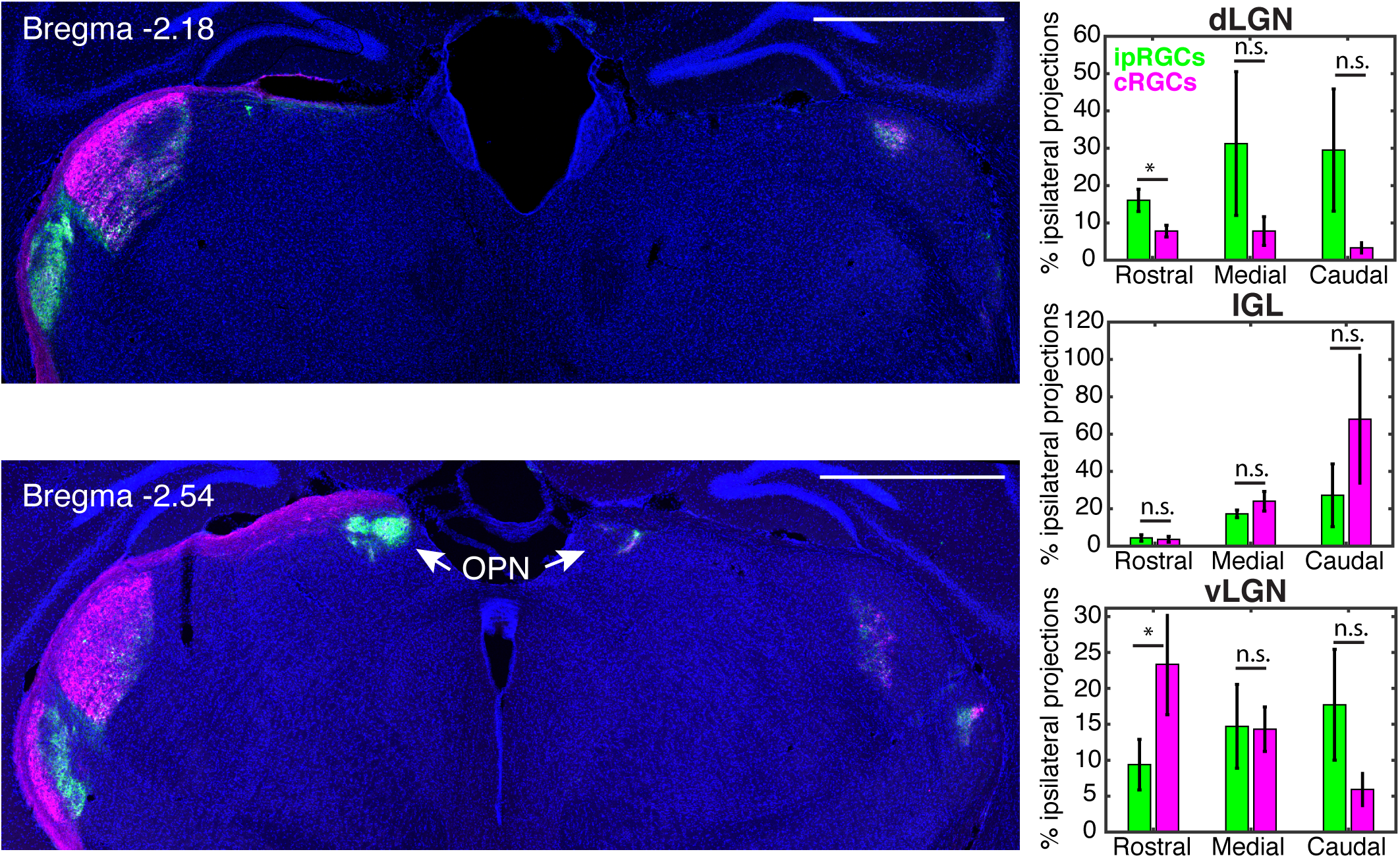
Ipsilateral ipRGC and conventional RGC projections to the geniculate complex. Coronal brain sections (enlarged from Figure 4) showing contralateral (left) and ipsilateral (right) innervation of ipRGCs (green) and conventional RGCs (magenta) to the rostral (Bregma -2.18) and caudal (Bregma -2.54) geniculate complex. The olivary pretectal nucleus (OPN) is indicated by arrows in the caudal brain section. Right side panels show the percent of ipRGC (green) and conventional RGC (cRGCs, magenta) innervation to the dorsal lateral geniculate nucleus (dLGN), the intergeniculate leaflet (IGL), and the ventral LGN (vLGN). Ipsilateral innervation between ipRGCs and conventional RGCs was not significantly different except in rostral dLGN and vLGN (Student’s paired t test, p < 0.5 indicated by * and p > 0.5 shown as n.s.)

### The caudal ipsilateral IGL receives conventional RGC innervation

The IGL has been implicated in circadian photoentrainment, and, more recently, in nonphotic behaviors such as sleep and feeding (Edelstein and Amir, 1999; Fernandez et al., 2020; Huang et al., 2019; Shi et al., 2020). Consistent with its role in circadian photoentrainment, it is not surprising that the contralateral IGL was predominately innervated by ipRGCs and received little to no conventional RGC innervation (Figure 4 and Figure 6). The ipsilateral IGL also received substantial ipRGC input (Figure 7). Surprisingly, in contrast to the dLGN, there was more ipsilateral input from conventional RGCs at the far caudal IGL (Figure 5 and Figure 7). The conventional RGC projections overlapped, however, completely with ipRGC input (Figure 7, lower panel).

**Figure 6.**
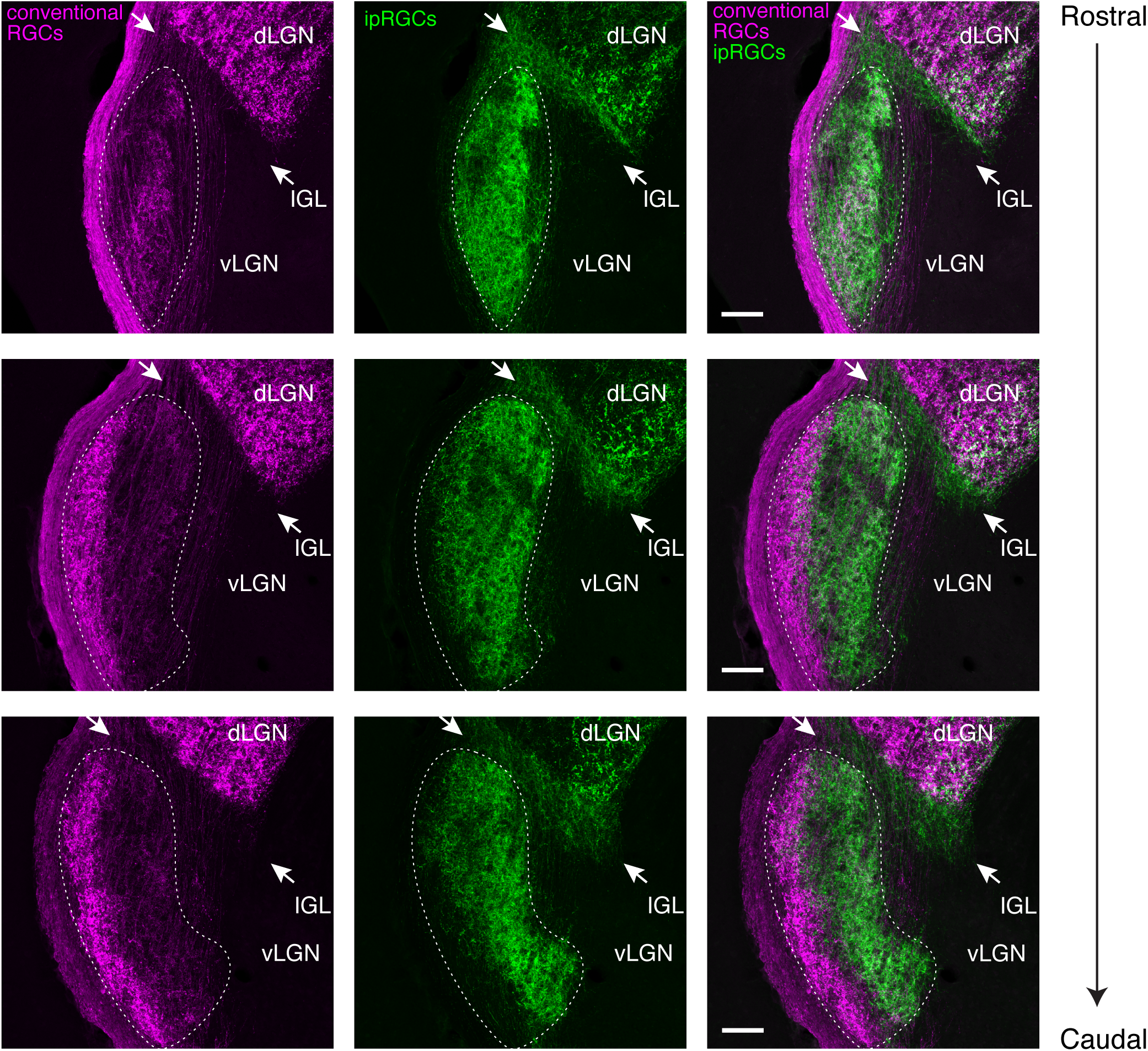
ipRGC and conventional RGC projections to the contralateral vLGN and IGL. Conventional RGC (magenta) and ipRGC (green) innervation to the contralateral ventral lateral geniculate nucleus (vLGN) and intergeniculate leaflet (IGL). Retinal innervation to the vLGN is outlined (dashed lines). The IGL is indicated by arrows between the vLGN and dorsal LGN (dLGN). Scale bars 100 µm.

**Figure 7.**
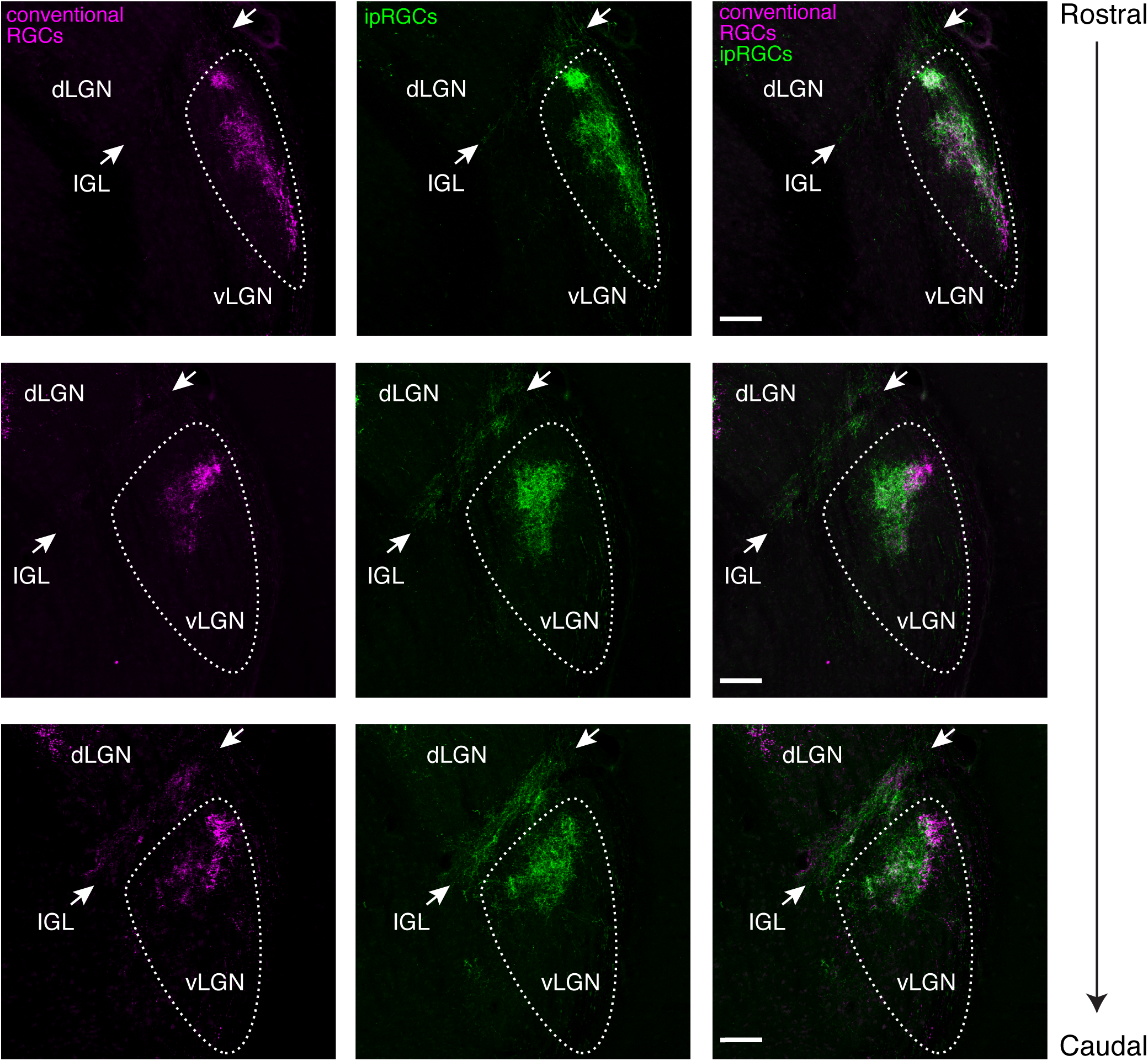
ipRGC and conventional RGC projections to the ipsilateral vLGN and IGL. Detailed view of conventional RGC (magenta) and ipRGC (green) innervation to the ipsilateral ventral lateral geniculate nucleus (vLGN) and intergeniculate leaflet (IGL), as seen in Figure 4. The vLGN is outlined (dashed lines). The IGL is indicated by arrows between the vLGN and dorsal LGN (dLGN). Scale bars 100 µm.

### ipRGC and conventional RGC innervations stratify in the vLGN

The vLGN is a retinorecipient region associated with non-image forming behaviors, although its function is not well understood (Davidowa and Albrecht, 1992; Harrington, 1997; Huang et al., 2019; Monavarfeshani et al., 2017; Shi et al., 2020). The vLGN also shows retinotopic responses to visual stimuli necessary for gaze control, suggesting a role in image-forming vision as well (Nagata and Hayashi, 1984). We observed a remarkable separation of ipRGC and conventional RGC projections within the caudal vLGN (Figure 4 and Figure 6). In the rostral vLGN, ipRGC and conventional RGC projections overlapped entirely (Figure 4 and Figure 6). In the caudal vLGN, conventional RGCs innervated only the lateral side of the vLGN, leaving the medial (core) side to ipRGC projections without any overlap (Figure 4 and Figure 6). In the ipsilateral vLGN, stratification was still visible with areas of overlap (Figure 7) and it has the characteristics of both the dLGN and IGL. Namely, in the rostral portion, more conventional RGCs innervate the ipsilateral vLGN, whereas, more ipRGCs innervate the ipsilateral vLGN in the caudal region (Figure 5). Together, these data show that there are clear stratification patterns in the lateral geniculate complex between ipRGCs and conventional RGCs.

### Conventional RGC and ipRGC projections overlap to innervate the OPN

The OPN shell is known to receive innervation from ipRGCs that are required for normal pupil constriction (Chen et al., 2011; Hattar et al., 2002, 2006). The OPN core has also been shown to receive ipRGC innervation (Chen et al., 2011; Delwig et al., 2016; Ecker et al., 2010; Quattrochi et al., 2018), although its function remains unknown. We found that conventional RGCs substantially innervated the core but avoided the shell of the contralateral OPN (Figure 8). In the ipsilateral OPN, projections from conventional RGCs and ipRGCs overlap (Figure 8).

**Figure 8.**
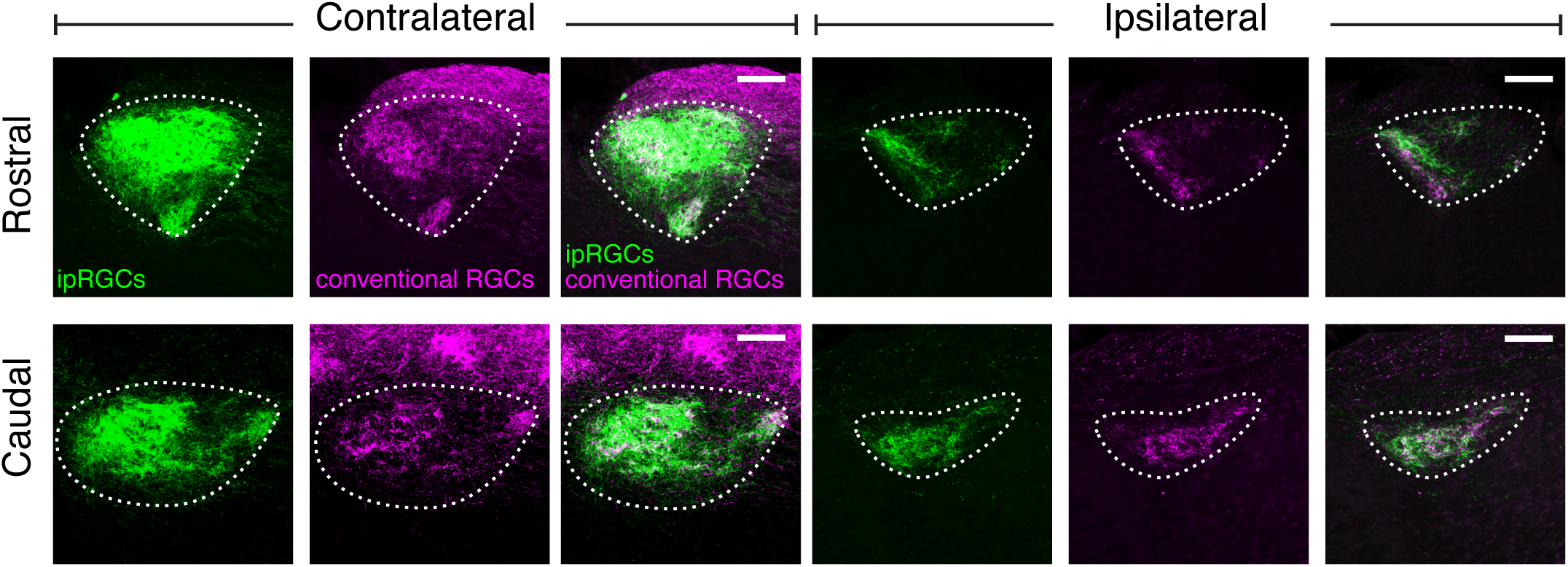
ipRGC and conventional RGC projections to the OPN. ipRGC (green) and conventional RGC (magenta) innervation to the contralateral and ipsilateral olivary pretectal nucleus (OPN). Retinal innervation to the OPN is outlined (dashed lines). Scale bars 100 µm.

### Retinorecipient targets with modest ipRGC innervation receive little input from conventional RGCs and weak ipsilateral innervation

The PHb mediates the direct effect of light on mood and is thought to be innervated primarily by ipRGCs (Fernandez et al., 2018). Similar to the projection patterns we observed in the SCN and vLGN core, we found that ipRGC innervations are the predominate retinal afferents to the PHb, with only a few conventional RGC puncta, indicative of minimal innervation (Figure 9). No ipsilateral innervation was visible in this region.

**Figure 9.**
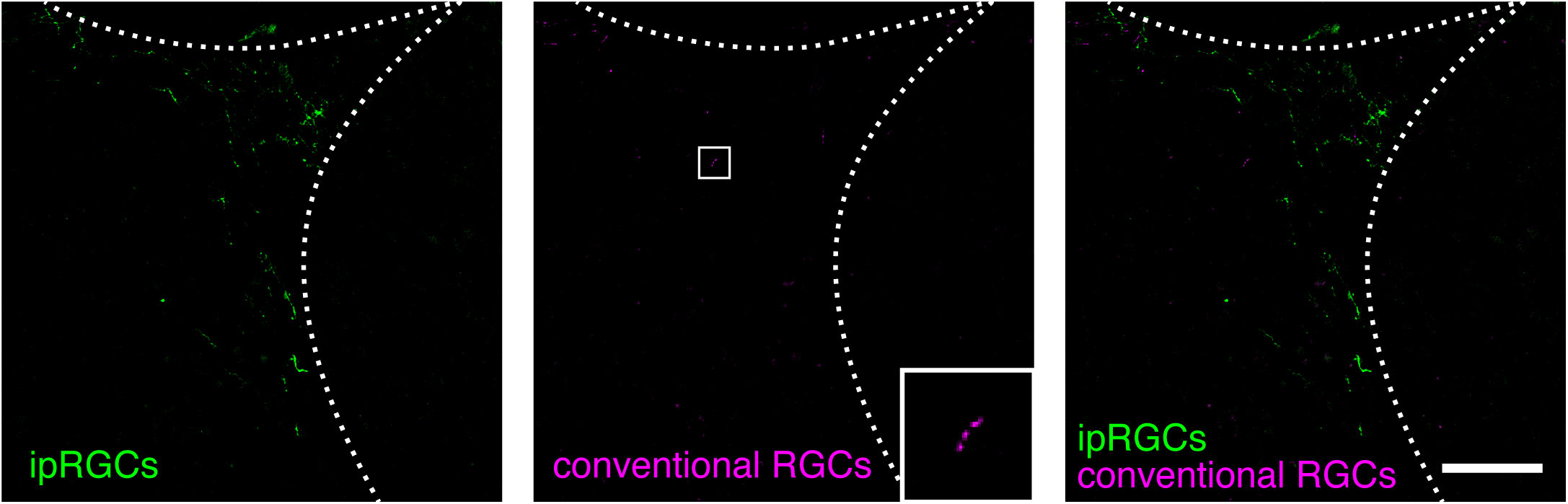
ipRGC and conventional RGC projections to the PHb. ipRGC (green) and conventional RGC (magenta, inset) innervation to the contralateral peri habenula (PHb). Dashed lines indicate the border between the hippocampus and the habenula. Scale bar 100 µm.

The ZI responds to some visual stimuli but its role in visual behaviors is not well understood (Wang et al., 2020; Zhao et al., 2019). At the border of the optic tract and ZI lies the peripeduncular nucleus, which we found is innervated exclusively by conventional RGCs (Figure 10). The significance of the retinal input to the peripeduncular nucleus is completely unknown, but it is likely innervated by a direction-selective RGC subtype (Rivlin-Etzion et al., 2011). The ZI, on the other hand, is innervated entirely by ipRGCs, except for a few conventional RGC fibers (Figure 10, inset). We did not observe retinal afferents in the ipsilateral ZI, as expected (Morin and Studholme, 2014).

**Figure 10.**
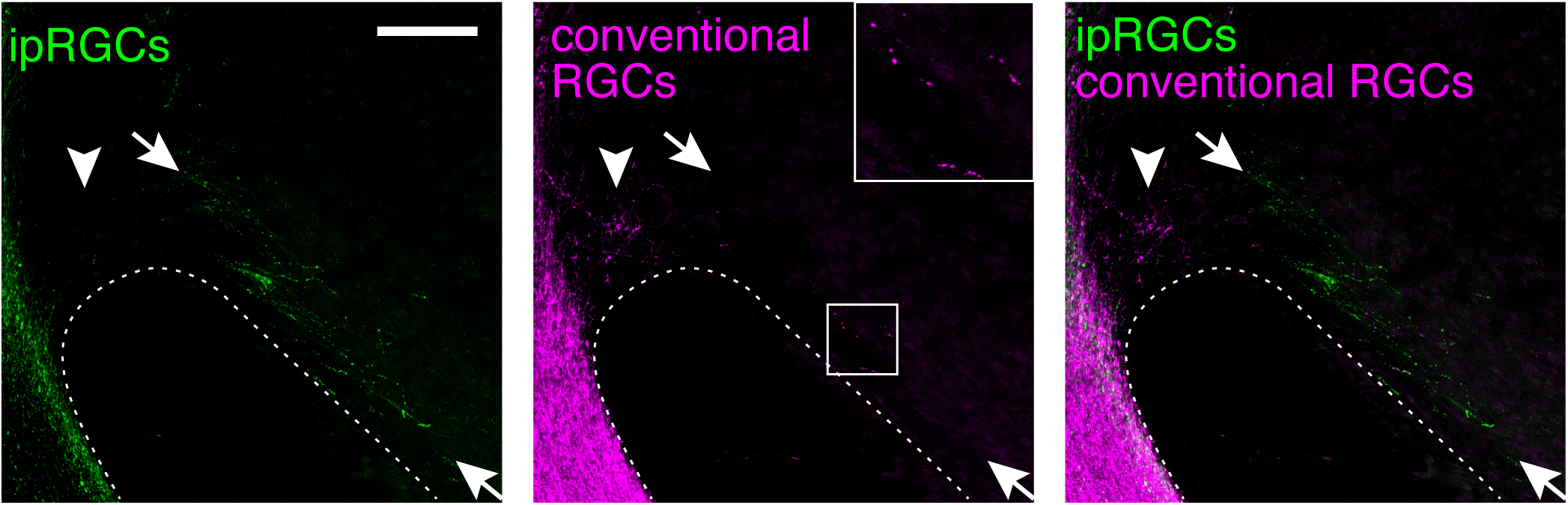
ipRGC and conventional RGC projections to the ZI. ipRGC (green) and conventional RGC (magenta, inset) innervation to the contralateral zona incerta (ZI). Arrowhead indicates retinal innervation to the peripeduncular nucleus. Retinal innervation to the zona incerta lies between the arrows. Dashed line outlines the cerebral peduncle. Scale bar 100 µm.

The SON is a neurotransmitter-rich nucleus whose visual function is unclear, although it has been implicated in metabolism and hydration, as well as lactation in nursing females (Cunningham et al., 2004). We found both conventional RGC and ipRGC innervation to the SON (Figure 11). Conventional RGC innervation was sparser than ipRGC innervation but covered the region almost as completely.

**Figure 11.**
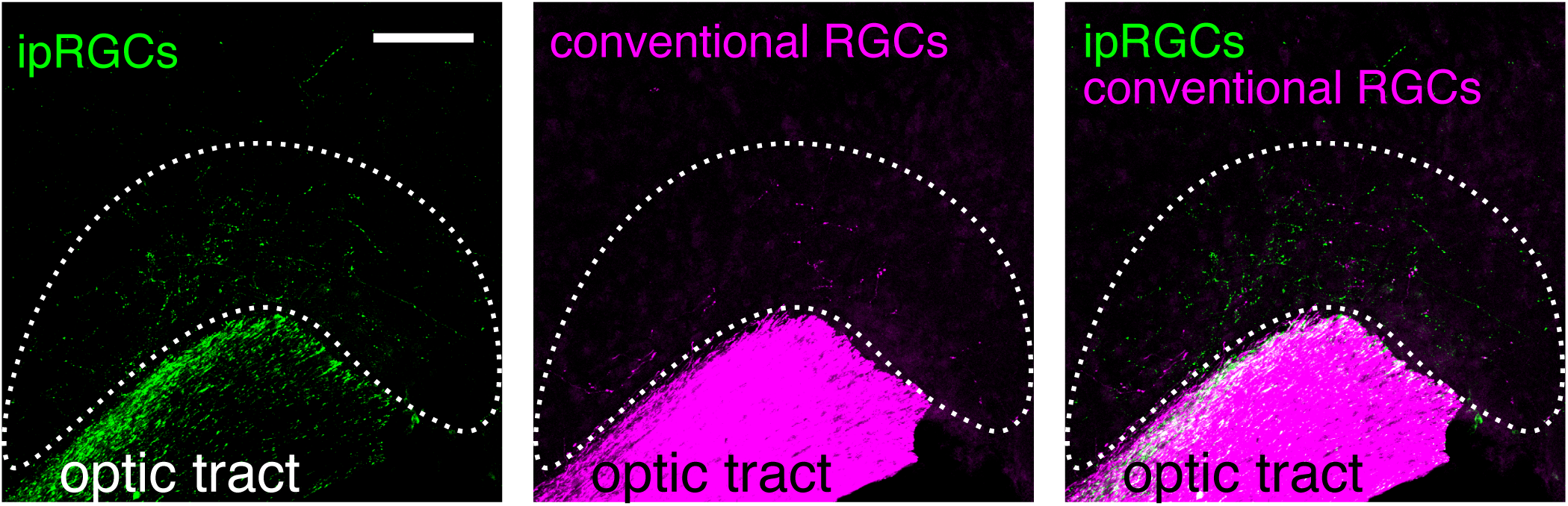
ipRGC and conventional RGC projections to the SON. ipRGC (green) and conventional RGC (magenta) innervation to the contralateral supraoptic nucleus (SON). Above the optic tract, retinal innervation to the SON is outlined (dashed line). Scale bar 100 µm.

## DISCUSSION

The role of ipRGCs in many non-image forming visual behaviors has been well-established, and their dense innervation to non-image forming brain nuclei is well known, but it remains unclear if ipRGCs are the sole conveyers of non-image forming visual information. Previous work has described ipRGC innervation patterns in comparison to all RGCs, but these studies could not readily delineate conventional RGC innervation to targets with dense ipRGC innervation due to the overlap in their labeling (Hattar et al., 2006; Morin and Studholme, 2014). Here, we used a dual recombinase labeling strategy to compare, in the same mouse, the innervation patterns of ipRGCs and conventional RGCs in a non-overlapping manner. We found that many non-image forming retinorecipient nuclei are innervated by conventional RGCs. The overlap and innervation patterns of ipRGC and conventional RGC projections vary drastically across retinorecipient nuclei and between ipsi and contralateral targets. This provides the intriguing possibility that conventional RGCs may modulate a subset of, but not all, non-image forming behaviors.

We were most surprised to find clear stratification and separation of ipRGC and conventional RGC inputs to the vLGN. Well-segregated cellular layers have been recognized in the dLGN and vLGN-like regions in primates and other diurnal mammals, but mice were not thought to have the same rigid organization in the geniculate complex (Kerschensteiner and Guido, 2017; Livingston and Mustari, 2000; Monavarfeshani et al., 2017). Some stratification of retinal subtype innervation to the mouse geniculate has been noted, but it is unclear if the stratification is indicative of non-overlapping retinal inputs (Kerschensteiner and Guido, 2017; Monavarfeshani et al., 2017). Our findings suggest that there is previously unrecognized organization in the vLGN that receives non-overlapping retinal innervation, consistent with recent data on cell type laminae within the vLGN (Sabbagh et al., 2020).

The pupillary light response relies on ipRGC innervation to the OPN shell majorly from M1 ipRGCs (Chen et al., 2011; Hattar et al., 2002, 2006). We found that the OPN receives overlapping ipRGC and conventional RGC innervation, except in the contralateral shell of the OPN, which receives only ipRGC input. The function of the OPN core is not well-understood (Dhande et al., 2018), but it is likely that ipRGCs and conventional RGCs both play a role in its visual response.

The IGL is another non-image forming nucleus that receives conventional RGC input. Excitingly, the innervation from conventional RGCs is limited to the caudal ipsilateral side (Figure 5, Student’s paired t test, p = 0.007). The IGL has been implicated in numerous non-image forming behaviors, making its conventional RGC innervation somewhat surprising (Edelstein and Amir, 1999; Fernandez et al., 2020; Huang et al., 2019; Shi et al., 2020). Furthermore, the increase in conventional RGC innervation to the IGL along the rostral-caudal and contra-ipsi axes fits well with previous studies suggesting underlying complexity within the cellular organization of the IGL (Moore and Card, 1994; Morin, 2013). It remains to be seen what role conventional RGCs will play in the ipsilateral IGL.

Removing ipRGC innervation to the SCN and PHb results in the loss of their light-modulated behavioral output (Fernandez et al., 2018; Güler et al., 2008; Hatori et al., 2008). Therefore, it is perhaps not surprising that ipRGCs were the predominant afferents to the SCN and PHb and it is unlikely that the few conventional RGC fibers we observed play any substantial functional role in these regions. Similarly, while the function of the ZI visual response remains to be elucidated, it is unlikely that conventional RGCs play a substantial role in its modulation due to their minimal innervation. The SON has been implicated in a myriad of behaviors (sleep and anesthesia, inflammatory pain processing, and thirst) but its role in vision and dependence on light modulation is unknown (Eliava et al., 2016; Jiang-Xie et al., 2019; Zimmerman et al., 2017). The SON received moderate conventional RGC input, especially when compared to the few puncta we see in the SCN, PHb and ZI. Our findings suggest that conventional RGC influence on the SON should not be ruled out.

The *Opn4*^*Cre*^ mouse labels all ipRGCs which are a heterogeneous population made up of at least 6 known subtypes. The Brn3b-negative M1 ipRGCs are known to exclusively innervate the SCN core (Chen et al., 2011). It is possible, or even likely, that retinal innervation specificity exists amongst other ipRGC and conventional RGC subtypes. As genetic mouse lines and tools to differentiate individual RGC subtypes become ever more available, it will be important to understand if innervations by all RGC subtypes segregate or overlap to contribute to visual function.

Taken together, we show that non-image forming retinorecipient brain regions are often not exclusive to ipRGC innervation. Our findings provide a basis for future experiments that will reveal how both ipRGCs and conventional RGC together modulate non-image forming behaviors for normal function.

## Acknowledgements

This project was supported by funding from the National Institute of Mental Health project number MH002964 (S.H.). We would like to thank Hui Wang in the Hattar lab for her management of the *Opn4*^*Cre*^ mouse line.

## METHODS

### Animals

All mice were handled in accordance with guidelines of the Animal Care and Use Committees of the National Institute of Mental Health (NIMH). RC::FLTG mice (Jackson Laboratory, Stock # 026932, (Plummer et al., 2015)) were crossed with *Opn4*^*Cre*^ mice (Ecker et al., 2010) to produce *Opn4*^*Cre/+*^;*Rosa26*^*CAG-tdTomato-eGFP/+*^ mice for experiments. Mice were group housed in a temperature and humidity-controlled room under a 12-hour light-dark cycle. Food and water were available *ad libitum*.

### Viral intravitreal injections

Adult *Opn4*^*Cre/+*^;*Rosa26*^*CAG-tdTomato-eGFP/+*^ mice (8-10 weeks) were anesthetized with isoflurane. The virus (Addgene plasmid pAAV-hSyn-Flpo, # 60663, (Xue et al., 2014), packaged into AAV2 at the NINDS Viral Production Core Facility) was placed on a piece of Parafilm and loaded into a glass microcapillary (Fisher Scientific, Cat. No. 11714) pulled needle (Sutter Instruments, Model P-2000). The glass needle was used to puncture the sclera of the right eye and 1.5µL of virus was injected into the vitreous chamber (Drummond Science Nanoinject II, Cat. No. 3-000-204). After the injection, mice were recovered from anesthesia and placed back into their home cages.

### Immunohistochemistry

Mice were deeply anesthetized 4 weeks after intravitreal injection and perfused with phosphate-buffered saline (PBS) followed by 4% paraformaldehyde. Eyecups and brains were removed for post fixation in 4% paraformaldehyde (1 hour and overnight, respectively). Retinas were isolated and washed in 0.5% Triton X-100 in PBS for 15 minutes three times. Blocking solution, with 3% donkey serum (Vector Labs), 0.5% Triton X-100, in PBS was applied to retinas overnight with gentle agitation at 4° C. Retinas were washed and incubated with primary antibodies in 1% donkey serum, 0.5% Triton X-100, in PBS solution for 3 days at 4° C. Primary antibodies used were chicken anti-GFP (1:2000, Abcam, Cat. No. ab13970), goat anti-tdTomato (1:1000, LSBio, LS-C340696) and rabbit anti-melanopsin (1:1000, Advanced Target Systems, Cat. No. AB-N38). Retinas were washed and incubated with secondary antibodies overnight at 4° C. Secondary antibodies used were donkey anti-chicken 488nm (1:1000, Jackson ImmunoResearch, Cat. No. 703-545-155), donkey anti-goat 555nm (1:1000, Invitrogen, Cat. No. A-21432), and donkey anti-rabbit 647nm (1:1000, Invitrogen, Cat. No. A-31573). Retinas were washed and then mounted flat in Vectashield (Vector Labs, H-1000). Brains were sunk in 30% sucrose, frozen, and sliced into 40 µm think coronal sections on a cryostat (Leica). Brain sections were treated as retinas with the following exceptions. Sections were blocked in 3% goat serum for 2 hours at room temperature. Sections were incubated in primary antibodies overnight at 4° C. Primary antibodies used were chicken anti-GFP (1:2000, Abcam, Cat. No. ab13970) and rabbit anti-RFP (1:1000, MBL International, Cat. No. PM005). Sections were incubated in secondaries for 2 hours at room temperature. Secondary antibodies used were goat anti-chicken 488nm (Invitrogen, Cat. No. A-11039) and goat anti-rabbit 546nm (Invitrogen, Cat. No. A-11035). Sections were mounted in Flouromount-G with DAPI (ThermoFisher, Cat. No. 00-4959-52).

### Imaging and analysis

Fluorescent images of retinas and brain sections were taken on a confocal microscope (Nikon, Eclipse Ti2). Images in the manuscript represent the maximal projections of a Z-stack (3µm step size) taken through an entire 40 µm thick coronal brain section. Any contrast or brightness adjustments were applied to the entire image in Fiji. To determine the percent ipsilateral projections, first, the maximal projection images were binarized using the Otsu method in Fiji. Regions of interest were created based on DAPI staining around the SCN, dLGN, IGL and vLGN. The total area of the contralateral and ipsilateral ipRGC or conventional RGC innervation to each region of interest was measured in the same coronal brain slice. For the geniculate measurements, coronal brain slices were grouped intro rostral, medial, caudal groups (rostral distance from Bregma -2.00 to -2.18, medial distance from Bregma -2.30 to -2.46, caudal distance from Bregma -2.54 to -2.70, Bregma based on (Franklin and Paxinos, 2007), n = 4-6 brain slices per group). The percent ipsilateral innervation was calculated by ipsilateral area over total retinal innervation multiplied by 100 to give percentage.

